# Ecology, phylogeny, and the evolution of developmental duration in birds

**DOI:** 10.1101/797498

**Authors:** Christopher R. Cooney, Catherine Sheard, Andrew D. Clark, Susan D. Healy, András Liker, Sally E. Street, Camille A. Troisi, Gavin H. Thomas, Tamás Székely, Nicola Hemmings, Alison E. Wright

## Abstract

The duration of the developmental period represents a fundamental axis of life history variation in animals, yet broad insights regarding the drivers of this diversity are currently lacking. Here using embryological data combined with information on incubation and fledging periods for 3096 species, we test key mechanistic and adaptive explanations for the evolutionary diversification of developmental durations in birds. First, using data on embryonic development for 20 model species, we show that developmental phases associated primarily with growth are longer and more variable than earlier phases, consistent with a role for allometric constraint in determining the duration of development. Second, using phylogenetic comparative methods, we find that avian developmental durations retain a strong imprint of deep evolutionary history, and that after accounting for these effects, body size differences among species explain less variation (5-22%) in developmental period lengths than previously thought. Finally, by collecting data for a suite of potential explanatory variables, our analyses reveal broad-scale ecological correlates of developmental durations, including variables associated with the relative safety of the developmental environment (e.g. nest height, insularity) and pressures of breeding phenology (e.g. migration). Overall, our results reveal that the combined effects of species’ body size, ecology, and phylogenetic history can account for 62-93% of the variation in developmental durations across birds, providing broad-scale quantitative insight into the relative importance of mechanistic constraints, adaptive evolution and evolutionary constraints in shaping the diversification of this key life-history trait.

## INTRODUCTION

A fundamental goal in ecology and evolution is to explain the vast diversity of life-history strategies observed in nature (1-3). The duration of the developmental period represents a fundamental axis of life history variation (4) and varies from days to several years among animal species. Attempts to explain variation in developmental duration across species typically fall into two broad categories (5). A first set of hypotheses, focusing on the role of mechanistic constraints, predict that developmental periods vary among species largely as a result of negative scaling between mass-specific growth (metabolic) rates and body size (6-8). In other words, larger species grow relatively slower compared to smaller species and therefore take longer to develop. Alternatively, a second set of hypotheses emphasise the role of selection and adaptation in generating interspecific differences in developmental durations. These adaptive ideas stem from classic life-history evolution theory (2, 4, 9) and assume that the external context of the organism drives the evolutionary optimisation of growth rates (and hence developmental period), within the constraints imposed by size and other assumed trade-offs (5). However, despite both sets of hypotheses being rooted in robust theoretical arguments, confidence in each is undermined by a lack of broad-scale empirical support and uncertainty exists regarding the relative importance of different factors for explaining broad-scale variation in developmental rates, particularly after accounting for phylogenetic effects (10).

Here we address this problem by conducting a global-scale phylogenetic comparative analysis of avian developmental periods and other life history traits. Birds are particularly suited for such analyses because accurate information on the duration of major avian developmental periods (incubation and fledging) is available for many species, as well as for many relevant aspects of species’ biology, ecology and distribution. In addition, detailed data on embryonic developmental stages are available for several taxonomically-diverse bird species, permitting the integration of large-scale comparative analyses with fine-scale investigation into differences in species’ developmental rates. To quantify broad-scale variation in overall developmental period length across birds we follow previous studies [e.g. refs. (11-14)] and use the sum of incubation and fledging periods, thereby capturing variation in both pre-natal and post-natal development rates. Furthermore, by combining data on incubation and fledging duration, we are able to define a second variable, which we refer to as the *incubation fraction*. This variable captures differences among species in the balance between prenatal (incubation) and postnatal (fledging) development periods and is calculated as the duration of the incubation period divided by the total developmental duration (incubation + fledging). This is useful because in birds, as in other animals, the proportion of prenatal to postnatal development varies among species, raising important questions regarding the factors explaining these differences (e.g. developmental mode, predation risk, etc.) (15, 16). However, while the importance of such factors have been examined in some taxa (17-27), their importance is rarely tested at broad phylogenetic scales.

To test key mechanistic and adaptive explanations for the evolutionary diversification of developmental durations in birds, we collected data from two different sources. First, we extracted standardised estimates of developmental timepoints from a taxonomically-diverse sample of 20 bird species with existing information on embryonic development. We predicted that if mechanistic constraints related to growth rates play an important role in determining avian developmental durations, then developmental phases associated primarily with growth should be longer and more variable across species than earlier phases concerned mostly with cell differentiation and body plan formation. Second, we compiled information on incubation and fledging durations for a total of 3096 bird species from 176 families and 39 orders, and combined this in a phylogenetic comparative framework with comprehensive data for a suite of variables that have previously been linked to avian developmental durations (17-27). Specifically, we tested variables related to species’ body mass, life history, parental care, nesting behaviour, ecology, ambient climate and biogeography. This two-scale approach—combining detailed observations of embryonic development with a broader comparative dataset—allows us to (i) to identify the phase(s) of avian development contributing most to inter-specific differences in developmental duration, (ii) investigate the strength of phylogenetic signal in trait values, and (iii) directly test and compare the relative importance of multiple potential underlying factors for determining developmental period length. Specifically, we use this approach to quantify the relative roles of mechanistic constraints and species’ ecology in shaping the evolution of avian developmental durations at a global scale.

## RESULTS AND DISCUSSION

A central prediction of mechanistic explanations for variation in developmental durations is that bigger species take relatively longer to develop. This is because growth is fuelled by metabolism which scales negatively with body size, such that larger species have lower relative metabolic rates than smaller species, and thus take proportionally longer to develop (6-8). A corollary of this is that if mechanistic constraints related to growth rate and body size during ontogeny represent an important rate-limiting step in offspring development, then phases of development associated with growth should be more variable across species, and account for a greater proportion of total developmental time, than non-growth periods.

To test the prediction that growth periods represent the longest and most variable phases of offspring development, we conducted a fine-scale analysis of developmental rates in a taxonomically diverse set of species (*n* = 20) with existing information on the timing of key developmental stages (Fig. S1, Appendix S1). Specifically, we examined four distinct phases in avian ontogeny spanning both pre-hatching (incubation) and post-hatching (fledging) periods (Fig. 1A). Phases 1 and 2, defined on the basis of Hamburger-Hamilton (HH) stages 1-24 and 25-32, respectively, correspond to periods of chick embryogenesis. These early stages of prenatal development consist primarily of cell differentiation and embryo formation rather than absolute growth (28), and we therefore consider these as “non-growth” phases. In contrast, phases 3 and 4, corresponding to HH33 to hatching and hatching through to fledging, are primarily concerned with periods of prenatal (phase 3) and postnatal (phase 4) growth of existing structures (see Supplementary Material for extended justification of these developmental phases). We used this framework to investigate variation in the duration and partitioning of avian offspring development.

**Fig. 1.**
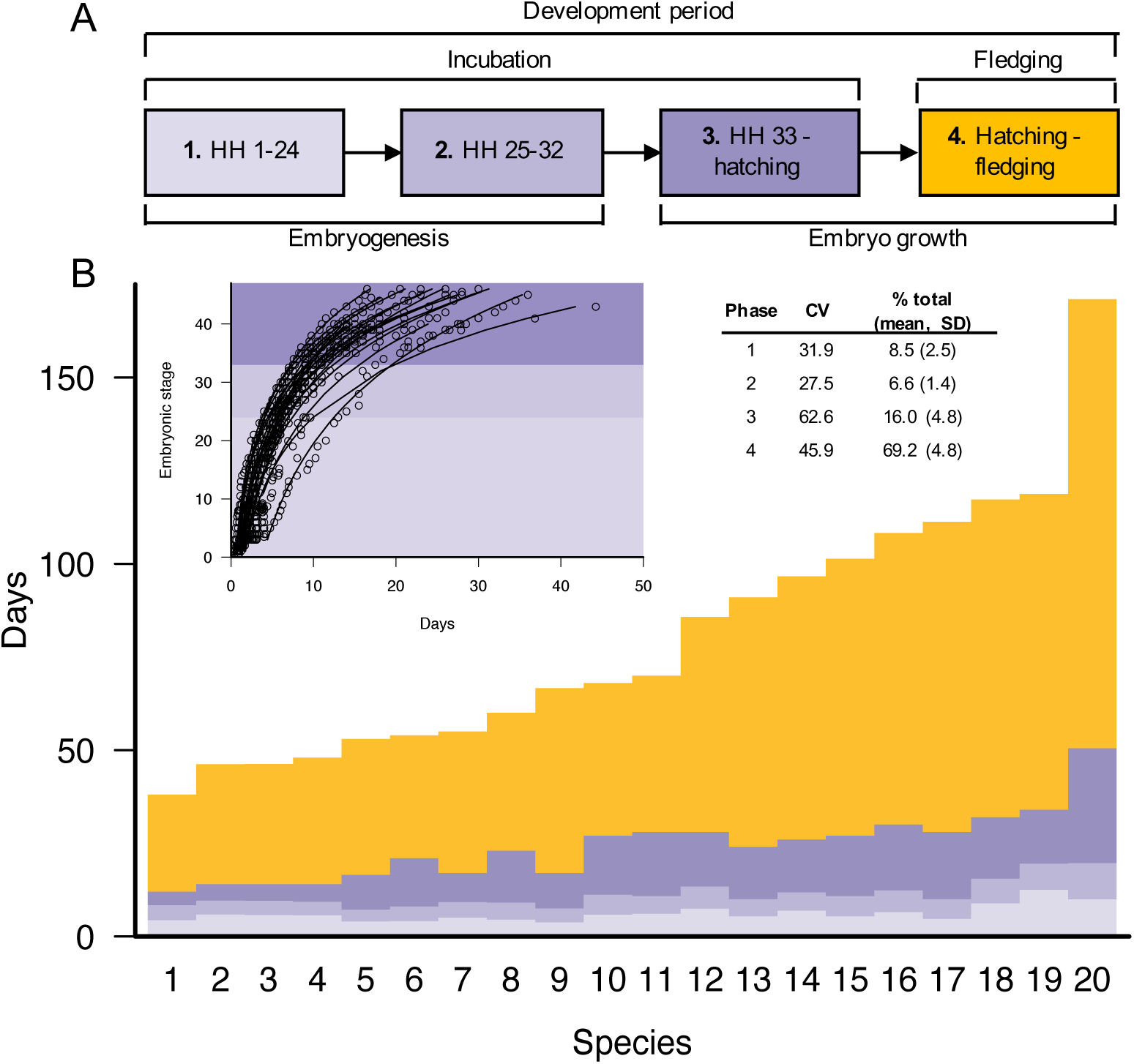
The duration of avian developmental phases. A, schematic illustrating four distinct phases of avian ontogeny. Phase 1 and phase 2 corresponding to Hamburger-Hamilton stages (HH) 1-24 and 25-32, respectively, represent embryonic developmental stages primarily associated with embryogenesis (i.e. non-growth). In contrast, phases 3 (HH33 to hatching) and 4 (post-hatching fledging period) correspond to developmental periods consisting largely of growth. B, stacked bar chart showing time intervals associated with phases 1 to 4 for 20 bird species for which information on the timing of embryonic developmental stages was available, with species are ordered by total developmental duration. Species codes are shown in Fig. S1. Inset graph shows the staging data and fitted curves used to estimate the time points separating phases 1-3. Inset table reports the coefficient of variation (CV) and percentage of total developmental period length (% total) accounted for by each of the four phases.

We found that the durations of developmental stages associated with embryogenesis (phases 1 and 2) account for only a small proportion of the variance in overall developmental duration across species (Fig. 1B). At these early stages of development, all bird embryos – regardless of species identity and eventual adult body size – are of comparatively similar size and therefore expected to have approximately similar growth rates (Fig. 1B). In contrast, the durations of growth phases (phases 3 and 4) are longer and more variable than non-growth phases and account for a far greater proportion of the variance in developmental duration among species (Fig. 1B). The longer duration of growth phases relative to embryogenesis phases is consistent with the well-characterised phenomenon of declining growth rates over ontogeny, caused by decreases in the ratio of energy acquisition to energy loss as developing organisms increase in size (5). Furthermore, greater variance in the duration of growth phases relative to non-growth phases (as indicated by coefficient of variation scores; Fig. 1B) is also consistent with greater size-related effects on the later stages of development. As development progresses, offspring of different species become increasingly different in size and therefore exhibit far greater disparity in relative growth rates compared to earlier stages of development.

Our observation that the developmental phases associated primarily with growth are longer and more variable than earlier non-growth phases predicts that body size should explain a significant amount of variation in developmental durations due to metabolic scaling rules. To test this idea more broadly, we collected data on developmental durations (incubation and fledging periods) for 3096 bird species covering the breadth of the avian phylogeny (Fig. 2A). In our dataset, overall developmental durations ranged from ∼20 days in some passerine species (e.g. *Volatinia jacarina*) to >350 days in some seabird lineages (e.g. *Diomedea*). Likewise, estimates of the proportion of development allocated to incubation relative to fledging (i.e. the incubation fraction) also varied markedly across species, ranging from ∼0.15 (e.g. *Struthio camelus*) to >0.95 in certain landfowl and shorebird species (e.g. *Megapodius pritchardii, Synthliboramphus wumizusume*) (Appendix S1).

**Fig. 2.**
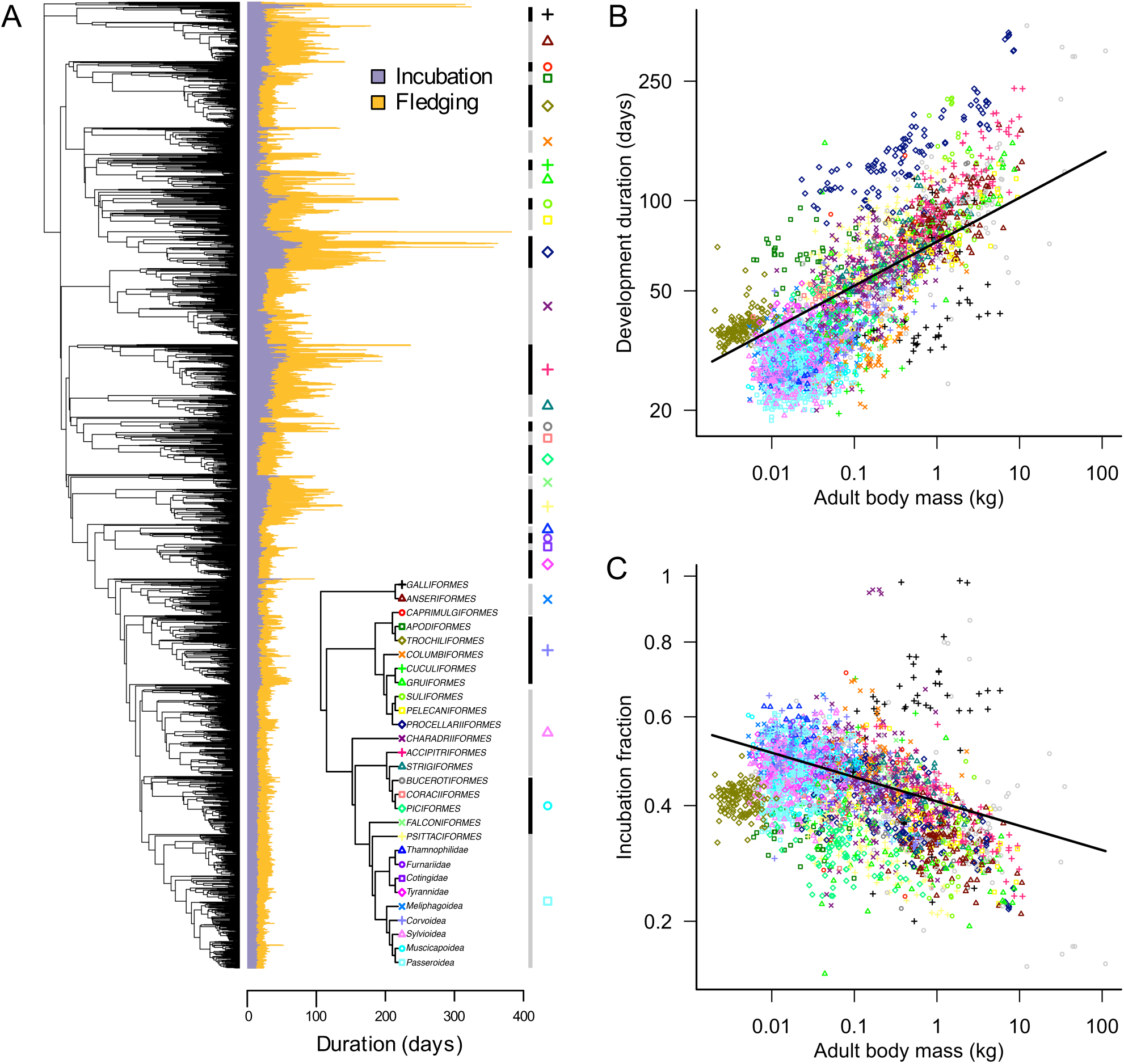
The diversity, phylogenetic distribution and allometry of development periods in birds. A, The phylogenetic distribution of incubation, fledging and overall development (incubation + fledging) period across 3096 species of birds. Inset tree schematic indicates the relationships among major taxonomic groups (>20 spp.) and provides a key for the plotting symbols used throughout the figure. B and C, allometric relationships of (log-transformed) development period length (B) and (square root-transformed) incubation fraction (C) with (log-transformed) adult body mass. Lines indicate the regression lines estimated by phylogenetically-controlled regression.

Before addressing relationships with body size, we first quantified the extent of phylogenetic signal in avian developmental durations. Phylogenetic signal measures the degree to which variation in species’ trait values covaries with phylogenetic relatedness and when estimated using Pagel’s λ model (29) varies on a scale from 0 (species trait values are unrelated to phylogeny) to 1 (species trait values follow Brownian motion expectations). Fitting Pagel’s λ model to our dataset, we found that developmental variables exhibited strong phylogenetic signal, with λ values [95% CI] of 0.93 [0.91, 0.94] and 0.86 [0.83, 0.89] for developmental duration and incubation fraction, respectively. This reflects a pervasive pattern in our dataset, that species within clades tend to exhibit similar developmental durations and incubation fractions (Fig. 2A), such that on average closely related species have more similar trait values than more distantly related species.

Against this backdrop, we used phylogenetic regression (30) and variance partitioning techniques (31) to test the relationships between body size and avian developmental duration, and to compare the contributions of predictor variables (body size) and variance components (phylogenetic effects) to the overall fit of the model. Using this approach, as predicted we found that overall developmental duration is positively related to body size across bird species (Fig. 2B). Furthermore, for incubation fraction we found that the offspring of larger-bodied species have proportionally shorter incubation periods relative to fledging periods (Fig. 2C), presumably reflecting energetic and/or ecological constraints associated with laying and/or developing in larger eggs. In both cases, we used adult body mass values as our index of body size across species, which we consider to represent a useful albeit imperfect proxy for offspring size at the end of development. However, we note that results were similar when we use an alternative proxy for offspring size (initial egg mass) (Appendix S1). Variance partitioning revealed that the partial *R*^2^ values associated with these phylogenetically-adjusted allometric relationships were 0.22 and 0.05 for developmental duration and incubation fraction, respectively. In contrast, the partial *R*^2^ values associated with the phylogenetic (covariance) components of each model were far greater: 0.79 and 0.59, respectively.

The significant relationships we observe between body size and developmental durations are in line with our predictions based on embryological data and provide broad empirical support for the role of size-related constraints in determining both the duration and partitioning of avian developmental periods (6-8). However, after accounting for phylogenetic effects, we found that the importance of body size for explaining variation in developmental durations across birds was surprisingly low, particularly considering that early tests implied that as much as 85% of inter-specific variation in incubation period could be explained by body size effects (32-34). In contrast, our comparatively low estimates for the variance explained by body size (5-22%) support the conclusion that, while important, allometric constraints play a more minor role in determining the length and partitioning of avian developmental periods than once thought (10). Instead, our quantitative estimates indicate that a greater proportion of the variance in avian developmental durations is attributable to phylogenetic history rather than body size. This finding is apparent in the observation that species within clades typically share similar developmental duration values that are largely unrelated to variation in body size both within and between clades (Fig. 2 and Fig. S2).

The existence of substantial mass-independent differences in developmental periods among bird lineages is intriguing, as it raises questions regarding the relative importance of mechanistic constraints versus adaptation in generating interspecific diversity in avian developmental periods. However, such questions have yet to addressed at broad scales (10). The idea that selection and adaptation play an important role in driving the evolution of developmental periods is rooted in classical life-history optimisation theory, the central tenet of which is that species adapt their life-history strategies to maximise fitness within particular ecological contexts (2, 4, 9). A range of factors have been suggested to be important in driving the evolution of developmental duration, many of which relate to either (i) constraints imposed by environmental or ecological limits to the resources available for reproduction, or (ii) selection imposed by increased mortality of parents and/or offspring. Field studies focusing on one or a few species have provided critical insight into relationships between ecology, selection and variation in developmental periods (17-27), but the restricted nature of these studies, combined with their often-conflicting results, have made broad conclusions difficult to draw. To test the relative importance of adaptation in explaining broad-scale variation in developmental periods, we studied the individual and combined effects of 16 variables related to behavioural, ecological, environmental, and life-history variation across species (plus body size) that have previously been linked to patterns of selection acting on avian developmental periods (17-27) (Fig. 3).

**Fig. 3.**
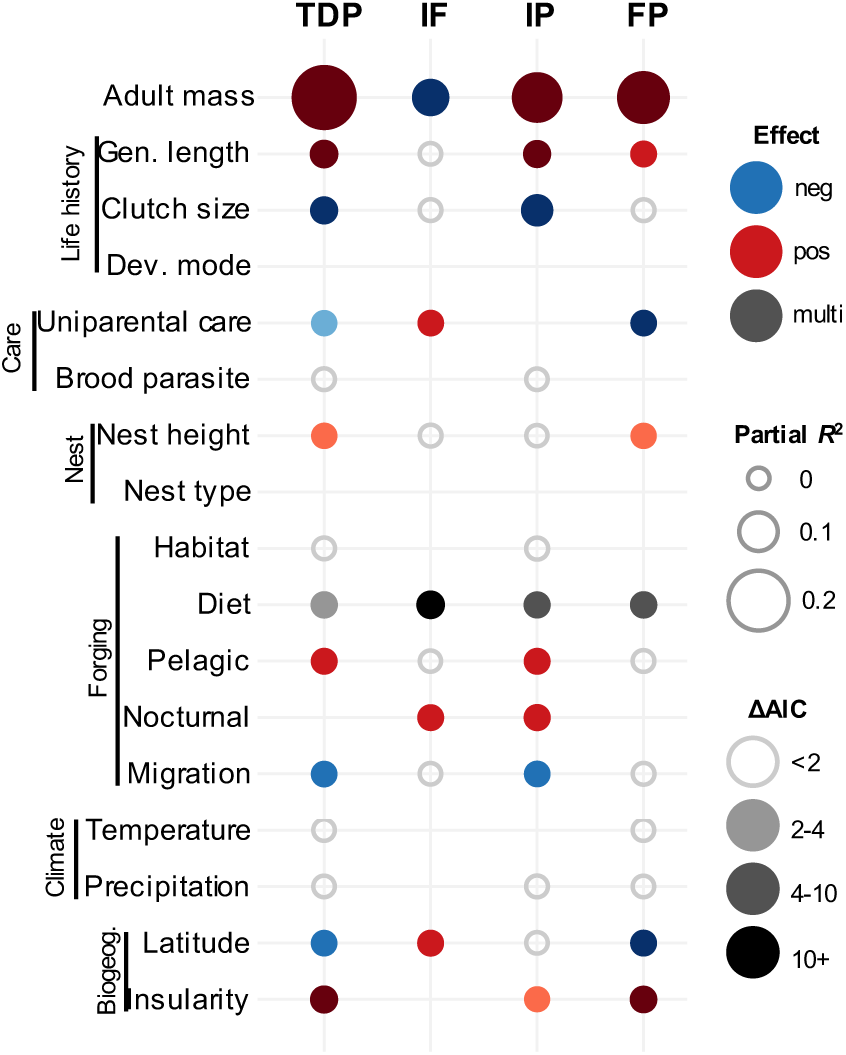
Predictors of the duration and partitioning of developmental period lengths in birds. A, Phylogenetically-controlled multi-predictor models of total developmental period (TDP), incubation fraction (IF), incubation period (IP) and fledging period (FD). Unfilled circles indicate factors that were significant as single predictors but not significant in a multi-predictor model. Gaps indicate factors that were not significant (ΔAIC > 2) as single predictors as single predictors and were therefore not included in the multi-predictor model. Note: factors with filled grey points (e.g. Diet) represent categorical variables with >2 (‘multi’) levels. ΔAIC values indicate the change in model support when the focal predictor was dropped from the model, with larger ΔAIC values indicating greater support for the importance of a predictor. Sample sizes (number of species) for the models were 1665, 1685, 1935, 1665 for TDP, IF, IP, and FP, respectively.

Our analyses revealed several important correlates of variation in avian developmental durations. First, after testing each predictor separately (see Fig. S3-6), we found strong relationships between several variables and developmental duration and incubation fraction across species (Appendix S1). By combining all significant single predictors in multi-predictor models, we were then able to identify sets of important predictors with unique effects that are independent of phylogeny. We found that, in addition to being larger, species with longer overall developmental durations tend to be longer lived, with smaller clutches, bi-parental care, elevated nest heights, vertebrate-eating/scavenging dietary niches, and pelagic foraging ecologies (Fig. 3; Appendix S1). These species also tend to be non-migratory and have more equatorial and insular breeding-range distributions. For incubation fraction (Fig. 3A; Appendix S1), in addition to the negative relationship with body size, we found that species with proportionally longer incubation periods tend to have uniparental parental care, are typically insectivorous and nocturnal, and have more polar breeding-range distributions. In both cases, broadly similar effects were found using initial egg mass as an alternative proxy for body size (Fig. S7; Appendix S1). Partial *R*^2^ values for these models indicated that, after controlling for phylogenetic and body size effects, the unique effects of ‘ecological’ variables included in multi-predictor models accounted for ∼12% and ∼4% of the variance in developmental duration and incubation fraction, respectively (Table 1). Interestingly, the magnitude of these effects were similar to those associated with body size (Table 1), implying that ecological and allometric effects (as measured here) explain roughly equivalent proportions of variation in developmental durations among bird species. Nonetheless, the variance associated with phylogenetic components indicated that phylogenetic effects remained a dominant source of variation in these models (Table 1). In total, these models incorporating body size, ecological, and phylogenetic effects accounted for 62-93% of the variation in developmental durations across species.

**Table 1.** (Partial) *R*^2^ values for model components explaining variation in avian developmental durations. Partial *R*^2^(individual components) and *R*^2^(full model) values are derived from phylogenetic multi-predictor models of total development period (TDP), incubation fraction (IF), incubation period (IP) and fledging (FP). Sample sizes and predictor sets are the same as those given in Fig. 3.

| Model component | TDP | IF | IP | FP |
| --- | --- | --- | --- | --- |
| Body size | 0.18 | 0.05 | 0.12 | 0.13 |
| 'Ecology' | 0.12 | 0.04 | 0.11 | 0.06 |
| Phylogeny | 0.62 | 0.43 | 0.70 | 0.52 |
| Full model | 0.91 | 0.62 | 0.93 | 0.84 |

These results have several important implications. Most notably, they show that behavioural and ecological variation among species can explain a significant amount of variation in developmental durations across species, consistent with an important role for selection in driving the evolution of avian developmental durations (4). In particular, three main ‘ecological syndromes’ appear to be associated with variation in developmental durations. First, longer developmental durations are generally associated with factors that presumably increase the safety of the developmental environment from predation threat or other mortality risks, such as nesting in relatively inaccessible sites (nest height), on islands (insularity), or having more than one parent to provide for and protect the offspring (biparental care). The idea that nesting in safe places may relax selection for rapid development is consistent with work by Remeš & Martin [ref. (21)], who found nestling growth rates to be positively associated with predation rates across passerines. Second, factors linked to phenological effects, such as breeding at temperate latitudes, insectivory, and migratory ecology, tend to be associated with shorter developmental durations. In species where reproductive success is driven largely by an individual’s ability to coincide their reproduction with peak seasonal food availability (4, 35, 36), the need to operate within a tight timeframe to avoid phenological mismatch is likely to select for rapid development (37). Third, several of the patterns we observe are also consistent with the importance of trade-offs between reproduction and survival for determining variation in avian developmental strategies. For example, shorter developmental periods in species with short lifespans and large clutches are consistent with selection for ‘fast’ life histories and greater investment in reproduction (independent of body size) (38, 39), while longer developmental periods among species with vertebrate hunting/scavenging diets are potentially explained by selection for slower development to mitigate costs associated with limited and/or unpredictable food availability (40, 41).

Furthermore, by considering predictors of incubation and fledging period separately, our results provide further insight into the patterns of selection generating underlying divergence in overall developmental duration and pre- versus postnatal allocation (Fig. 3, Appendix S1). For instance, our finding that nocturnal species have larger incubation fractions than diurnal species is seemingly driven by nocturnal species having relatively long incubation periods rather than particularly short fledging stages. This makes sense if lower daytime parental activity disproportionally reduces nest predation risk during the incubation period relative to the fledging period (26), thus relaxing selection for rapid development inside the egg. Similarly, the longer developmental periods of pelagic species are largely driven by relatively long incubation periods, which may be a consequence of selection for advanced development at hatching (22) or lower rates of egg predation due to inaccessible breeding locations (39). In contrast, our results show that species with uniparental care tend to have overall shorter developmental durations (and greater incubation fractions) largely because of reduced fledging durations. This is consistent with predictions for evolutionary associations between single parent care and short post-hatching offspring development periods (4, 42), but the direction of causality remains unclear. On the one hand, uniparental care may generate selection for rapid post-hatching offspring development to reduce the burden of care, but on the other hand short post-hatching periods may facilitate desertion by one of the parents (typically the male), implying a reversal in the direction of cause and effect (43).

Finally, our results challenge several assumptions regarding relationships between developmental durations and other factors at broad scales. In particular, ambient climate is predicted to shape broad-scale patterns of developmental rates in birds via its effect on egg temperature and parental behaviour (10). However, after controlling for the effect of other factors, we found no evidence that variation in environmental conditions (temperature and precipitation) was related to developmental duration across species. This finding supports the view that offspring are to a large extent buffered from variation in ambient environmental conditions by parental adaptations such as nest design, incubation efficiency, and provisioning rate (24, 44, 45). Surprisingly, we also found no significant relationships with developmental mode (precocial, semi-precocial, altricial) or nest type (cavity, closed, open, mixed). This is despite strong expectations for significant associations (15, 46-48) and seemingly large differences between groups in the raw data (see Fig. S3-6). We attribute these negative results to the effect of correcting for phylogenetic non-independence among species in our models. Variation in developmental mode and nest type have likely independently evolved only a limited number of times across the avian phylogeny (49) and so their effect on developmental durations cannot be disentangled from underlying patterns of shared evolutionary history and/or ecology (31). Thus, although developmental mode and nest type appear to co-vary with developmental durations, the generally assumed view that precocial species have longer developmental durations and greater incubation fractions (i.e. prenatal developmental) than altricial species is not supported when phylogenetic effects are taken into account. Greater clarity on whether factors such as developmental mode directly influence the evolution of developmental durations or are simply associated at broad scales via phylogenetically conserved constraints will likely come from integrating data on equivalent traits from other groups (e.g. all vertebrates) to generate sufficient independent phylogenetic replication to conclusively test these relationships.

Overall, our study reveals key drivers of developmental durations across the breadth of the avian phylogeny, providing broad, quantitative insight into the relative importance of mechanistic constraints and ecologically-mediated selection in explaining variation in key life-history traits. Furthermore, our results highlight the pervasive impact of phylogenetic history in shaping variation in species’ developmental durations. The close association between developmental duration, species’ traits, and phylogeny implies a strong signal of evolutionary conservatism, both in terms of species’ developmental durations and the combinations of factors (‘syndromes’) that co-evolve with them, echoing the conclusions from other large-scale phylogenetic analyses of avian life history traits (39, 49). While birds provide sufficient evolutionary replication to investigate the importance of many factors, phylogenetic constraints and evolutionary conservatism makes it difficult tease apart the effects of other, less labile, traits. Thus, a potentially fruitful avenue of future research would be to address these questions over even broader phylogenetic scales in order to better address the effects of body size, species’ traits, and phylogenetic constraints (e.g. mutation rates) on the evolutionary diversification of developmental durations.

## MATERIALS AND METHODS

### Data

We collected information on the timing of embryonic development for 20 species using data available in the primary literature (see Appendix S1 for references). For each species with existing data, we extracted information on the time taken for embryos to reach sequential stages of the Hamburger-Hamilton (28) scale, which represents a standard approach for describing and comparing rates of embryonic development across bird species (50). To ensure consistent measurements across species, one of us (NH) re-staged embryo development using data provided in the original publication. In cases where an alternative staging approach was used, we re-staged embryo development according to the Hamburger and Hamilton (28) scale using detailed descriptions and photographs provided in the original publication. In cases where a range of time points were reported for reaching a given stage, we used the average.

We collected information on prenatal (incubation) and postnatal (fledging) period lengths (days) for 3096 bird species listed in Appendix S1 from ref. (51) and major ornithological reference works (52). Following ref. (51), we define incubation period as the time (in days) between when the egg is laid and when it hatches, and fledging period as the time taken (in days after hatching) for offspring to be capable of flight (or for some species, leaving the nest). These variables have been used extensively in the comparative avian life history literature and represent standardised measurements of avian developmental periods that are broadly comparable across all bird species (51). Furthermore, the sum of incubation plus fledging time represents a commonly-used metric for species’ ‘total development period’ that integrates variation in both pre- and postnatal development times (11-14). While we acknowledge that in some bird lineages individuals continue to grow after fledging, we argue that in most cases post-fledging growth accounts for a relatively minor proportion of offspring development and as such the combined duration of incubation plus fledging periods represents an informative metric of the total development time.

To improve data quality we removed clear outliers that must reflect measurement error (i.e. incubation lengths < 8 or > 90 days; *n* = 6). In addition, we also assessed the extent of within-species variability in development period estimates (where available) relative to the extent of variation across all species by calculating repeatability (i.e. intra-class correlation) coefficients (53). Our dataset contained an average of 1.54 (range 1 to 9) measurements per species for incubation period and 1.47 (range 1 to 9) measurements per species for fledging period. Based on this data, estimated repeatability coefficients were 0.984 (95% CI = [0.982, 0.985]) for incubation period and 0.944 (95% CI = [0.939, 0.949]) for fledging period, implying low variability in estimates of developmental periods within species relative to variation between species. We therefore calculated mean values of incubation and fledging period per species (when multiple values were available) and from this calculated variables capturing total developmental duration (incubation + fledging) and incubation fraction [incubation / (incubation + fledging)].

Data on adult body mass (g), initial egg mass (g), generation length (days), clutch size, developmental mode (precocial, semi-precocial, altricial), parental care (uniparental, biparental), brood parasitism (parasite, non-parasite), minimum nest height (m), nest type (cavity, closed, open, mixed), habitat (forest dependency: high, medium, low, none), diet (omnivore, fruit/nectar, invertebrate, plant/seed, vertebrate/fish/scavenger), foraging (pelagic, non-pelagic), nocturnality (nocturnal, diurnal) and migration (sedentary, migratory) were extracted from standard avian trait databases (51, 54, 55) or scored directly from the literature [primarily ref. (52)]. Geographical variables, including temperature, precipitation, latitudinal midpoint, and insularity (continental, insular), were based on maps of species breeding distributions from http://www.datazone.birdlife.org (Version 9) combined with global climate (56) and landmass datasets (57). Further details of data compilation methods are given in the Supplementary Material. The full dataset, along with associated sample sizes, can be found in Appendix S1.

### Phylogeny

Our analyses are based on the taxonomy and phylogenies of ref. (58), which currently represent the only available ‘complete’ species-level phylogenetic hypothesis for all birds. To provide a phylogenetic framework for the species in our dataset (*n* = 3094), we downloaded 1000 ‘full’ trees (those containing all 9,993 species) from http://www.birdtree.org, which we then pruned to leave only the species represented in our dataset. We then used this tree distribution to generate a maximum clade credibility (MCC) tree, which provided the phylogenetic framework for our analyses.

### Analyses. *Staging data*

We categorized avian development into four discrete phases spanning both prenatal (embryonic; based on the descriptions of Hamburger & Hamilton [ref. (28)]) and postnatal (fledging) periods (Fig. 1A), and calculated the time taken for individuals to reach the end of each phase. Specifically, we estimated the time required for embryos to reach HH24 (phase 1), HH33 (phase 2), to hatch (phase 3) and finally to fledge (phase 4). To estimate the time points associated with reaching HH24 and HH33, we fitted curves of the form y = exp(a + b*x) using the R function ‘nls’ to describe the relationship between embryonic stage (x) and time (y) (Fig. S1). This allowed us to accurately infer time points associated with HH24 and HH33, even when such data were not explicitly reported in the original publication. Data on the later time points (hatching and fledging) were extracted either from the relevant staging paper directly or else imported from our broader comparative dataset.

### Allometric analyses

We tested the relationship between adult body size and developmental period variables using phylogenetic linear regression (30), controlling for phylogenetic relatedness by estimating Pagel’s lambda. For each variable, we also fit a model in which intercepts were estimated separately for major taxonomic groups (> 20 spp.) in order to generate mass-adjusted estimates of relative developmental durations (Fig. S2).

### Multi-predictor models

We used the same phylogenetic regression approach described above to test the relationship between predictor variables and variation in our developmental variables. First, we fitted individual (i.e. single) predictor models using all available data for each predictor. We then combined all individually important predictors into a multi-predictor model. In all cases, predictors were considered to be important if model support values dropped by >2 units (i.e. ΔAIC > 2) when the predictor was dropped from the model (59), with larger ΔAIC values indicating greater statistical support for the importance of a predictor. We checked for evidence of multi-collinearity among predictors in our multi-predictor models using variance inflation factors (VIF) and found no evidence of severe (VIF > 10) or even moderate (VIF > 5) multi-collinearity in any of our models (median VIF = 1.80; range = 1.01 – 4.13). *R*^2^ values for full models (including phylogenetic effects) and partial-*R*^2^ values associated with predictors were calculated using the ‘R2.lik’ function in the R package ‘rr2’ (31).

## ACKNOWLEDGEMENTS

We thank Kevin Laland for providing access to data, and Joseph Brown, Angela Chira, Yichen He, Emma Hughes and Jonathon Kennedy and XX reviewers for constructive comments on the manuscript. This work was funded by the European Research Council (grant number 615709 Project ‘ToLERates’), a Leverhulme Early Career Fellowship to CRC (ECF-2018-101), a Royal Society University Research Fellowship to GHT (UF120016), a NERC Independent Research Fellowship to AEW (NE/N013948/1), and a Royal Society Dorothy Hodgkin Research Fellowship to NH (DH160200).

## Supplementary material

### 1 Supplementary methods

#### 1.1 Extended justification for avian developmental phases

According to Hamburger and Hamilton (1951), stages 1-24 of embryo development (our ‘Phase 1’; Fig. 1A) are described exclusively in terms of embryogenesis – specifically the formation and organisation of the fundamental body plan. During this time (up to incubation day 4 in the chicken), changes in the embryo are characterised primarily by the number of somites, and then, once somites become difficult to see due to the development of the mesoderm, the development of limb-buds, visceral arches, and other externally visible structures. Similarly, stages 25-32 (incubation day 4-8 in the chicken; our ‘Phase 2’) are also characterised by rapid developmental changes in the wings, legs, and visceral arches, and can therefore also be considered as part of embryogenesis, as the differentiation of body structures (e.g. toes, mandible, etc.) is still ongoing.

In contrast, from stage 33 onwards, chick development is described primarily in terms of growth, rather than embryogenesis. Specifically, between stages 33-38 (incubation day 8-12 in the chicken), Hamburger and Hamilton describe changes in feather germs and eyelids to distinguish stages, both of which are already present in the developing embryo. Furthermore, from stage 38 (day 12 in the chicken) onwards, Hamburger and Hamilton explicitly state that no new structures are formed, and that chick development primarily comprises the growth of structures that already exist. Thus, from stage 38 onwards, Hamburger and Hamilton exclusively use measurements of beak and toe length (i.e. growth) to distinguish stages. We decided to include stage 33-38 into this ‘growth’ phase (our ‘Phase 3’) because although Hamburger and Hamilton were not exclusively using growth measurements to differentiate stages at this point, they were still using descriptions of growth based on existing structures only. Thus, we consider chick development from stage 33 to hatching (our ‘Phase 3’), and from hatching to fledging (our ‘Phase 4’), to constitute growth, in contrast to stages 1-32, which we consider to represent embryogenesis.

#### 1.2 Predictor variables

Data on mean adult body mass (g), egg mass (g), clutch size, diet (omnivore, fruit/nectar, invertebrate, plant/seed, vertebrate/fish/scavenger), foraging (pelagic, non-pelagic) and nocturnality (nocturnal, diurnal) were extracted directly from Wilman et al. (2014) and Myhrvold et al. (2015). We used the literature [primarily del Hoyo et al. (1992–2011) and Starck (1993)] to assign species to broad categories capturing variation in developmental mode (precocial, semi-precocial, altricial), parental care (uniparental, biparental), brood parasitism (parasite, non-parasite), nest type (cavity, closed, open, mixed). Nest height (m) was recorded as the (minimum) distance between the base of the egg cup and the ground for a given species reported in the literature. We extracted information on generation length (days), habitat (forest dependency: high, medium, low, none) and migration (sedentary, migratory) from http://www.datazone.birdlife.org following the approaches described in Cooney et al. (2018). Briefly, regarding species’ habitat classifications, in the BirdLife dataset species are assigned to one of four broad habitat categories, depending on whether they “do not normally occur in forests”, or exhibit “low”, “medium” or “high” levels of forest dependency. Similarly, BirdLife categorise species as “not a migrant”, “nomadic”, “altitudinal migrant” or “full migrant”. We converted this classification system into a binary variable capturing broad differences in species’ migratory tendencies, categorising each species as ‘non-migratory’ or ‘migratory’ (nomadic, altitudinal migrant or full migrant).

Variables relating to species’ geographical distributions are based on bird breeding range maps provided by BirdLife International and NatureServe (version 9; http://www.datazone.birdlife.org), rasterised to 1^°^ resolution. Following Jetz et al. (2008), we calculated average range-wide temperature and precipitation values for the warmest quarter (bio10 and bio18), extracted from the WorldClim2 database (Fick and Hijmans 2017), and we calculated species mean (absolute) breeding-range latitude values directly from grid cell occurrences.

Finally, insularity was determined by comparing species range maps to a dataset of global landmasses (GSHHG v2.3.6; http://www.soest.hawaii.edu/pwessel/gshhg/), and we defined insular species as those with >95% of their range occurring on islands as defined by Weigelt et al. (2013). Prior to analysis, incubation fraction was square-transformed, and the following variables were log-transformed: incubation, fledging and total developmental duration, adult body mass, generation length, clutch size and nest height. The full dataset, along with associated sample sizes, can be found in Appendix S1.

---

**Fig. S1.**
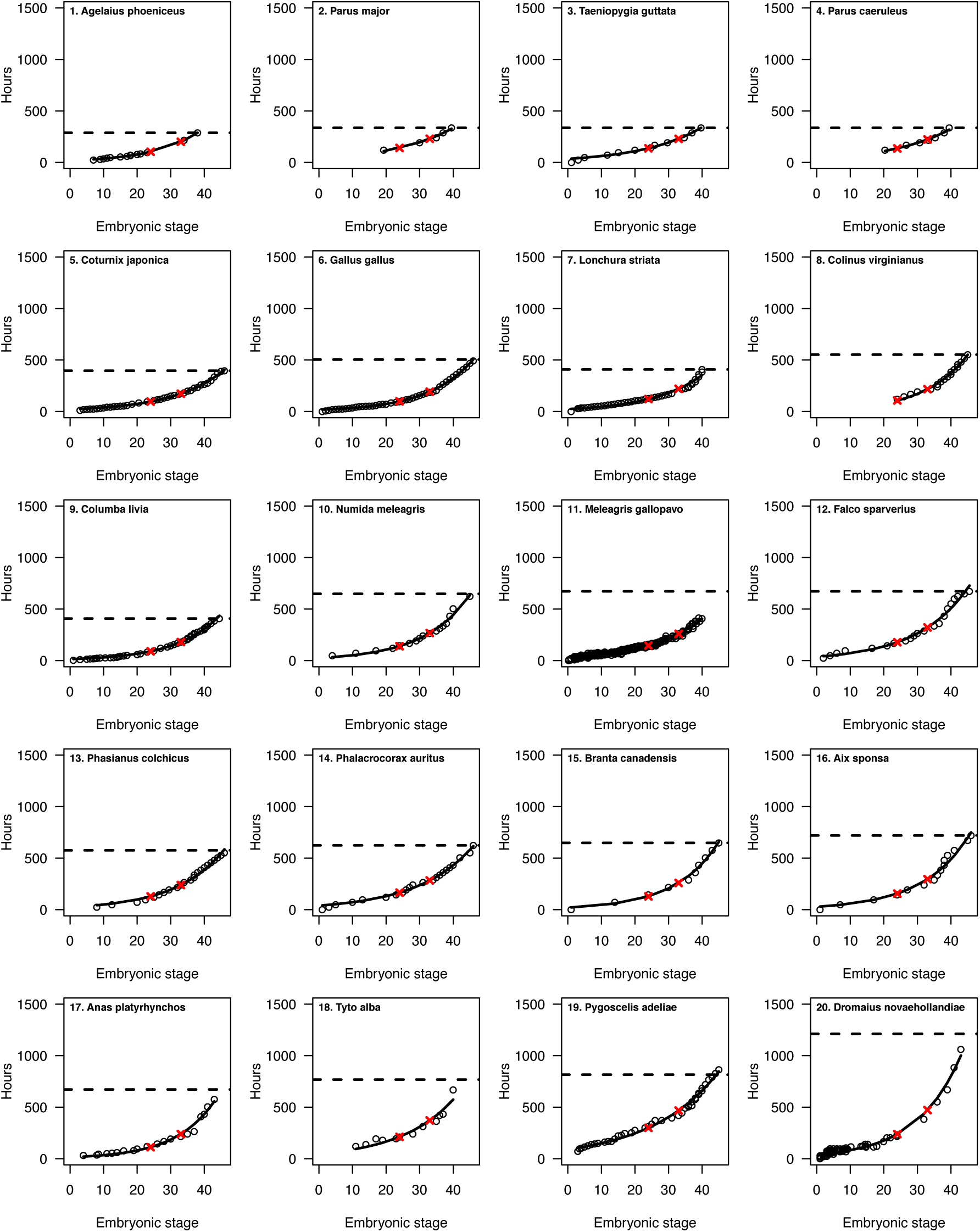
Individual embryonic development curves for 20 bird species. Points are observed data, and fitted lines come from fitting an equation of the form y = exp(a + b * x). Red crosses indicate the estimated time at which species reach embryonic stage 24 and 33, respectively. Dotted line indicates the hatching time, as reportedfromthe relevant literature.

**Fig. S2.**
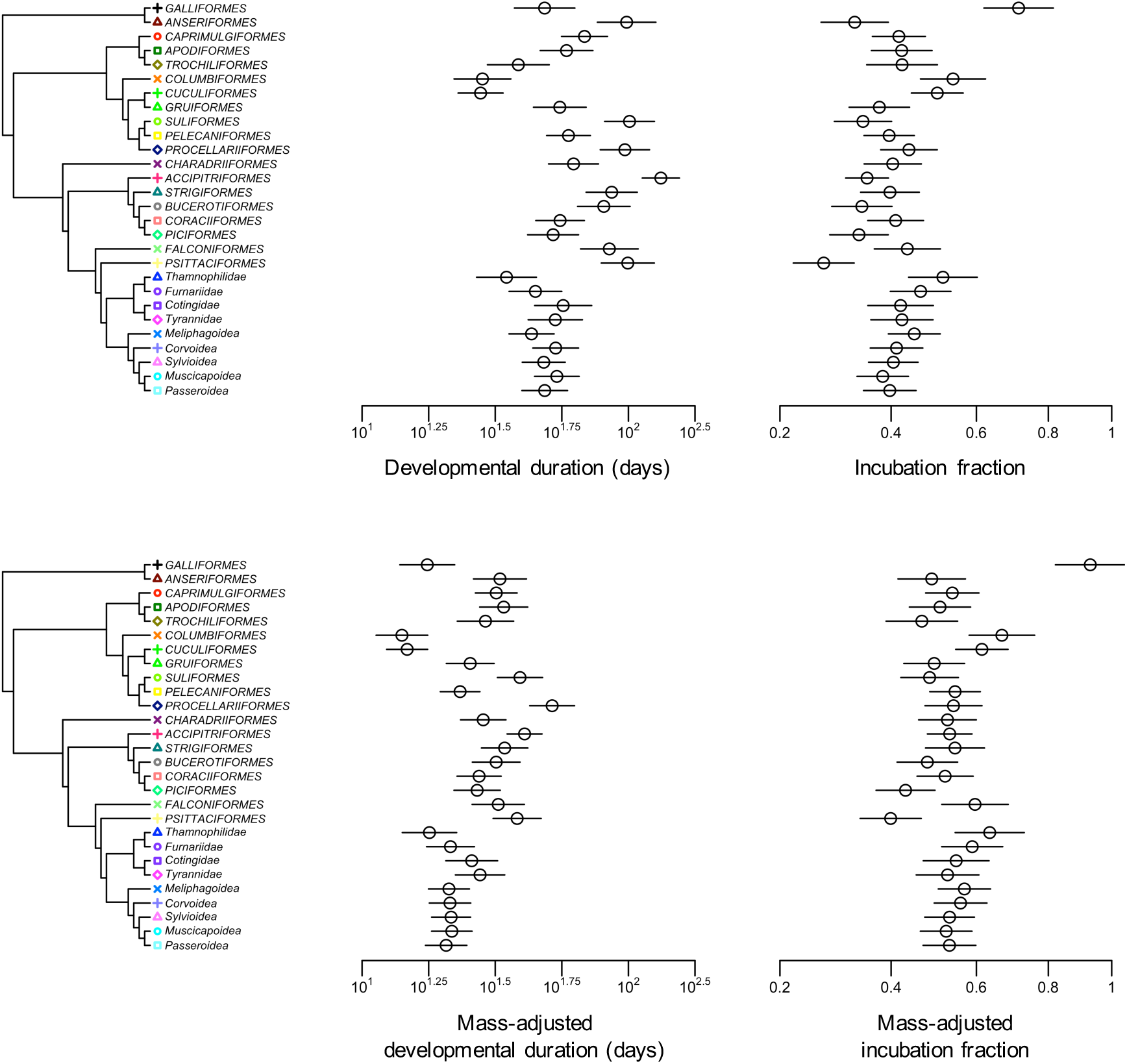
Variation in developmental durations of major avian clades. Relative developmental duration and incubation fraction values represent the *y*-intercepts from a model of the form *y* = *a* + *b* (log mass) in which major avian clades (>20 spp.) were permited to have unique intercepts (but parallel slopes). Horizontal lines indicate standard errors.

**Fig. S3.**
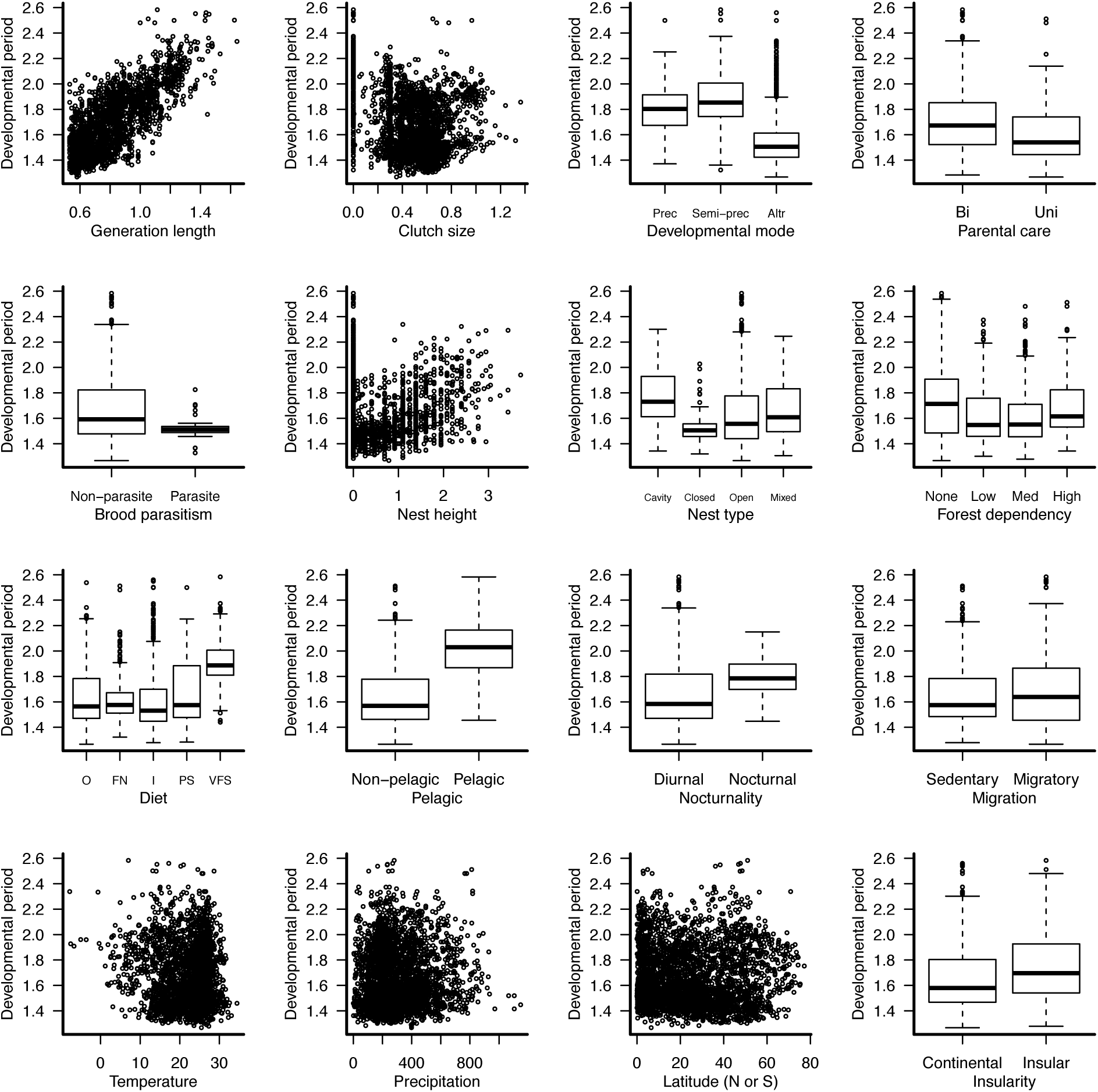
Relationships between (log_10_-transformed) total developmental period length and individual predictor variables. Sample sizes and regression statistics can be found in Appendix S1.

**Fig. S4.**
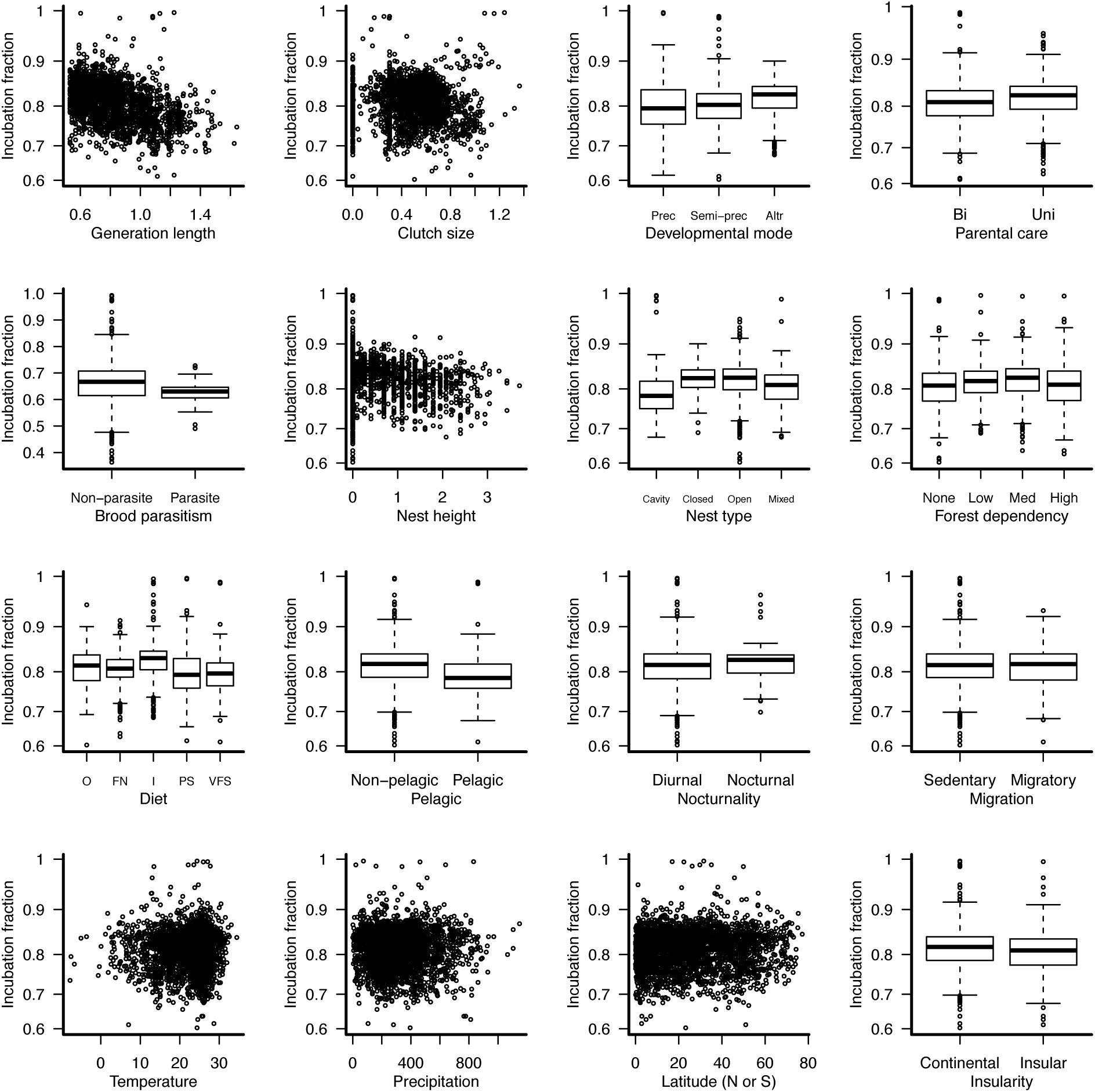
Relationships between (square root-transformed) incubation fraction and individual predictor variables. Samplesizes and regression statistics can be found in Appendix S1.

**Fig. S5.**
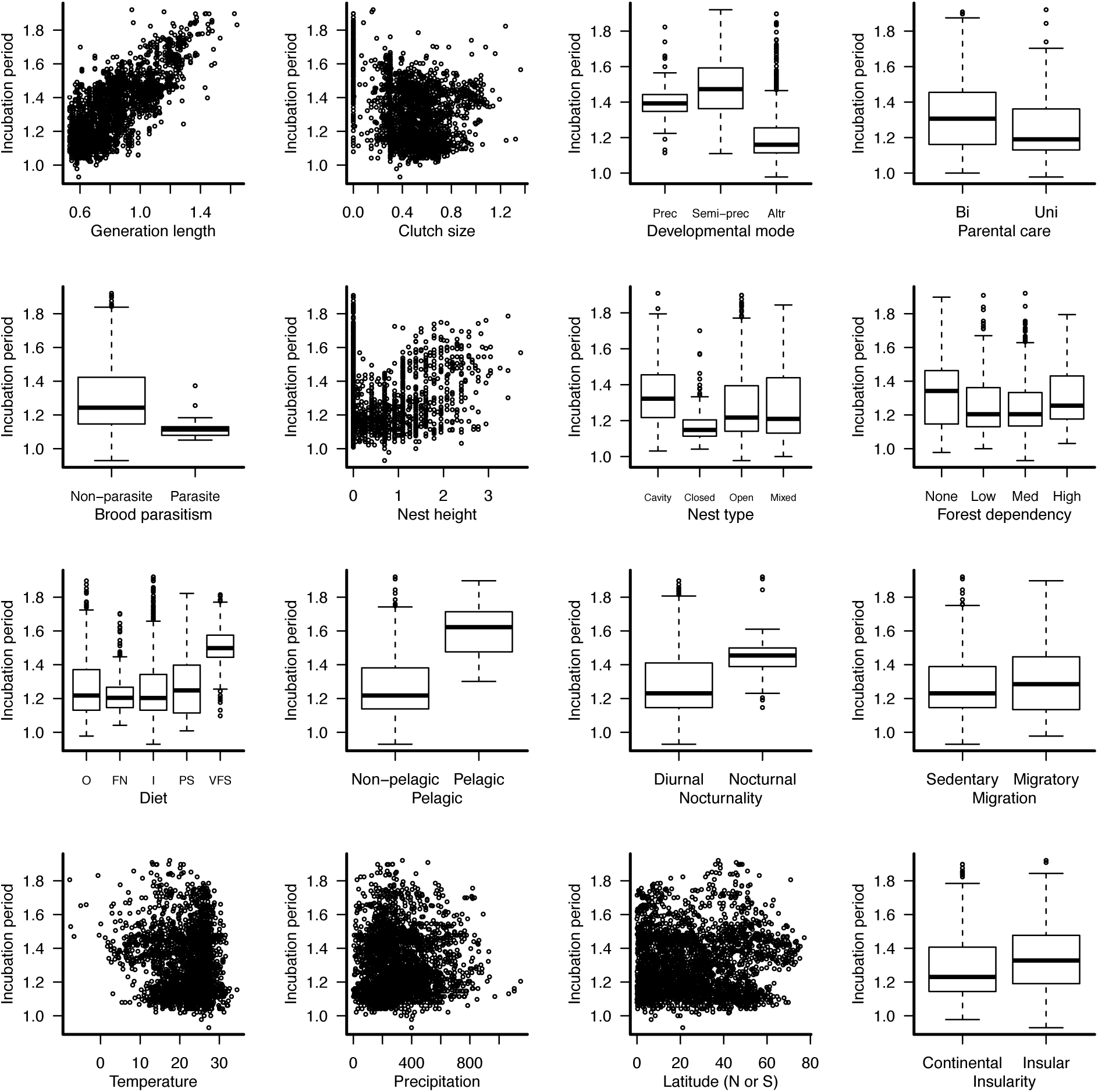
Relationships between (log_10_-transformed) incubation period length and individual predictor variables. Samplesizes and regression statistics can be found in Appendix S1.

**Fig. S6.**
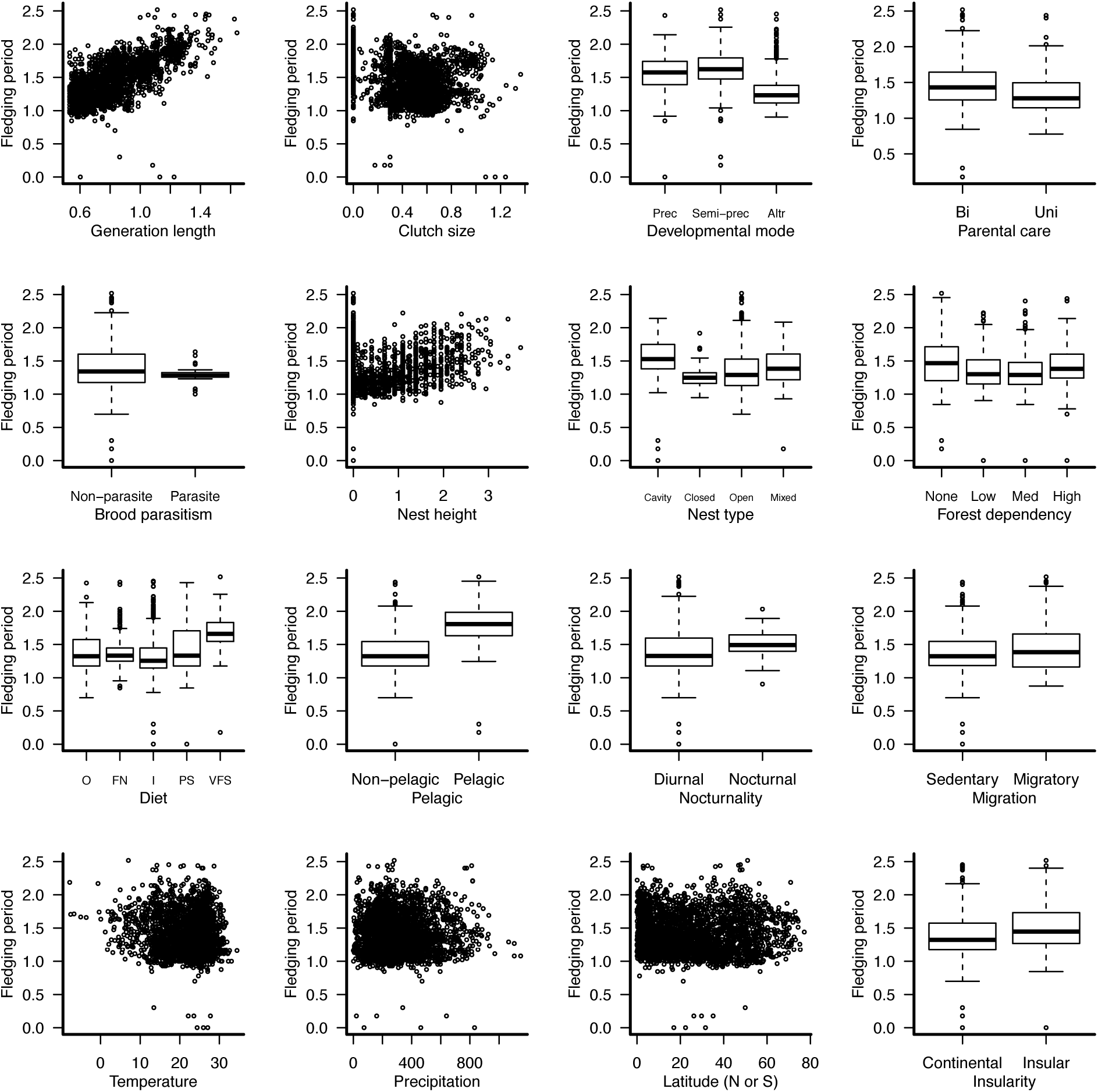
Relationships between (log_10_-transformed) fledging period length and individual predictor variables. Sample sizes and regression statistics can be found in Appendix S1.

**Fig. S7.**
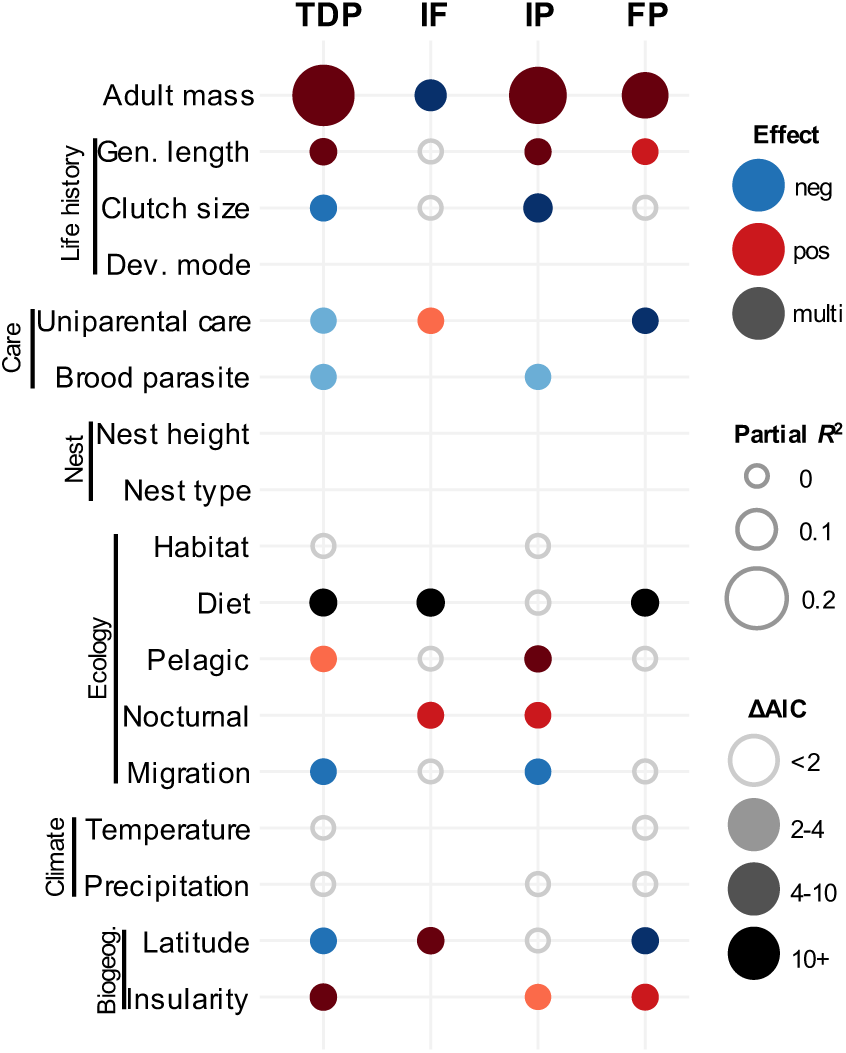
Predictors of the duration and partitioning of developmental period lengths in birds using egg mass as a proxy for (fledgling) body size. A, Multi-predictor models of total developmental period (TDP), incubation fraction (IF), incubation period (IP) and fledging period (FP). Unfilled circles indicate factors that were significant as single predictors but not significant in a multi-predictor model. Gaps indicate factors that were not significant (ΔAIC > 2) as single predictors as single predictors and were not included in the multi-predictor model. Note: factors with filled grey points (e.g. Diet) represent categorical variables with >2 (‘multi’) levels. ΔAIC values indicate the change in model support when the focal predictor was dropped from the model, with larger ΔAIC values indicating greater support for the importance of a predictor. Sample sizes (number of species) for the models were 2327, 1988, 1988, 2017 for TDP, IF, IP, and FP, respectively.

